# Shifting a Cellular Metabolic Landscape Identifies a Refractory Environment for Flavivirus Replication

**DOI:** 10.1101/2022.02.21.481365

**Authors:** Rebekah C. Gullberg, Nunya Chotiwan, M. Nurul Islam, Laura A. St Clair, Elena Lian, Thomas J. Edwards, Sudip Khadka, Christopher Teng, Barbara Graham, Kirsten Krieger, Amber Hopf-Jannasch, Douglas J. LaCount, John T. Belisle, Richard J. Kuhn, Rushika Perera

## Abstract

Host-targeted therapeutics to control viral infection are gaining prominence given the vulnerability of viral replication at select host-interaction points and the limited possibility of developing drug resistant mutants. Nevertheless, the chemical and biological impact of many host-targeted therapeutics on both the cell and virus has not been elucidated and remains a key complication. Previously, it has been demonstrated that inhibition of fatty acid metabolism has significant antiviral potential. Here, we use a multidisciplinary approach to demonstrate how inhibition of fatty acid biosynthesis creates a metabolically refractory environment that drives viral dependence on alternate metabolic pathways for survival. By profiling the global metabolic landscape following inhibition of fatty acid biosynthesis, we identified additional biochemical pathways that, when inhibited in combination with fatty acid biosynthesis, displayed increased antiviral potential. Our studies also demonstrated that there was a direct link between changes in cellular chemical composition and the ultrastructural membrane architecture induced by viral gene products. Utilizing inhibitors to change these metabolic environments significantly impacted early viral replication and disrupted the membrane architecture critical for the viral life cycle. Here, we have defined at a molecular level how shifting metabolic landscapes can be exploited to identify combinations of therapeutics that have a greater antiviral effect.

**Author Summary:** Dengue viruses are transmitted by *Aedes aegypti* mosquitoes which are prevalent in the tropical and subtropical regions of the world. These viruses cause over 350 million infections annually. There are no antivirals to combat infection and the only vaccine available is suboptimal. Since these viruses are obligate pathogens, they hijack lipid metabolic pathways in host cells to drive new lipid synthesis critically required for their replication. Mechanisms of how lipid synthesis impacts viral replication is unknown. These viruses also rearrange cellular membranes to form platforms for assembly of viral replication complexes. Here, for the first time, we show that virus-hijacking of *de novo* fatty acid biosynthesis pathways is required for the formation of membranous replication platforms and if inhibited disrupted synthesis of replicative form viral RNA. Importantly, these inhibitors drastically rearranged the metabolic landscape of the cell resulting in an activation of compensatory nucleotide synthesis pathways that allowed the virus to survive at a low level through the inhibition. However, if both pathways were inhibited in combination, infectious virus release was reduced to below detection limits. The study demonstrates how understanding the metabolic landscape altered by specific inhibitors can lead to the discovery of compensatory metabolic pathways and targets that in combination can enhance intervention efficacy.

## Introduction

With nearly 400 million annual infections worldwide, dengue viruses (DENVs) pose a significant global public health concern^1^. Infections with these viruses can be asymptomatic or range from dengue fever (DF), an acute and debilitating illness, to the more severe and life-threatening dengue hemorrhagic fever (DHF) and dengue shock syndrome (DSS). DENVs consist of four serotypes. Infection with one serotype does not cross-protect against infection with additional serotypes. In fact, secondary infections with different serotypes are associated with DHF/DSS^2^. There are no effective antivirals currently available to treat flavivirus infections, and the only vaccine on the market is sub-optimal^3–5^. Given the absolute reliance on host processes for successful viral replication, host-targeted therapeutics present an alternate strategy for rapidly re-purposing existing drugs as antivirals. In this study, we have explored the impact of therapeutics that alter host metabolism as an avenue to identify refractory metabolic landscapes and combination therapeutics that significantly alter viral replication and host pathology.

Electron microscopy analyses have indicated that the membranes of the endoplasmic reticulum (ER) form unique structures during infection and that these structures support viral replication^6, 7^. These membranes serve as the site of viral genome translation and polyprotein processing and function as a platform to concentrate substrates required for RC formation. They also protect viral replication intermediates from immune detection^8^.

The rearrangement of ER membranes requires a combination of cellular and viral proteins as well as changes in chemical composition of lipids to support the curvature and fluidity needed to form and maintain the viral RCs. Dengue virus serotype 2 (DENV2)-induced changes in the chemical composition of the ER and whole cell lipidome have been observed in mosquito cells as well as whole mosquitos^9, 10^. Changes to cellular lipid composition and concentrations are likely accomplished through viral control of cellular lipid synthesis enzymes. For example, fatty acid synthase (FAS) is a key lipogenesis enzyme that catalyzes the formation of palmitic acid (C16:0) from acetyl-CoA, malonyl-CoA and NADPH^11^. Inhibition of FAS is well-recognized to limit the genomic replication of numerous flaviviruses in multiple cell types^10, 12–16^. In DENV2-infected human cells, FAS was re-localized to sites of viral RNA replication^14^. Furthermore, viral protein NS3 was shown to interact with the dehydratase (DH) domain of FAS^11, 14^. NS3 also enhanced the activity of FAS *in vitro*^14^. FAS has a variety of commercially available inhibitors and is a high-profile therapeutic target for many other diseases including multiple cancers^17^, metabolic syndrome^18^ and nonalcoholic fatty liver disease^19^. Orlistat (Tetrahydrolipstatin) is an FDA-approved anti-obesity compound for its inhibition of gastric and pancreatic lipase^20, 21^. Orlistat covalently binds the active site serine in the thioesterase domain of FAS and prevents release of palmitic acid (C16:0). As a lipase inhibitor, Orlistat prevents hydrolysis of triglycerides from dietary sources and thus, thwarts absorption of fat as free fatty acids (FAs)^22, 23^. C75 is another FAS inhibitor which has a different mechanism of FAS inhibition (Fig 1A). While Orlistat prevents the release of the final product of FAS, palmitic acid (C16:0) and blocks reuse of the enzyme^22, 23^, C75 impedes FA chain elongation and prevents malonyl-CoA from binding the ketoacyl synthase (KS) domain of FAS^24^. This causes accumulation and/or re-direction of malonyl-CoA from fatty acid biosynthesis to its other functions such as interfering with the β-oxidation of fatty acids^25^. Here, we utilized Orlistat as an example to dissect the consequences of drug treatment on the metabolic landscape of human cells and the mechanisms driving DENV2 inhibition. We hypothesized that re-purposing therapeutics available for metabolic diseases as antivirals would be effective given the viral dependence on these metabolic pathways for their lifecycle. We also hypothesized that by examining the shifting metabolic landscapes influenced by inhibitors such as Orlistat we could identify additional inhibitors that can work in combination for a more potent reduction of viral replication.

**Fig 1.**
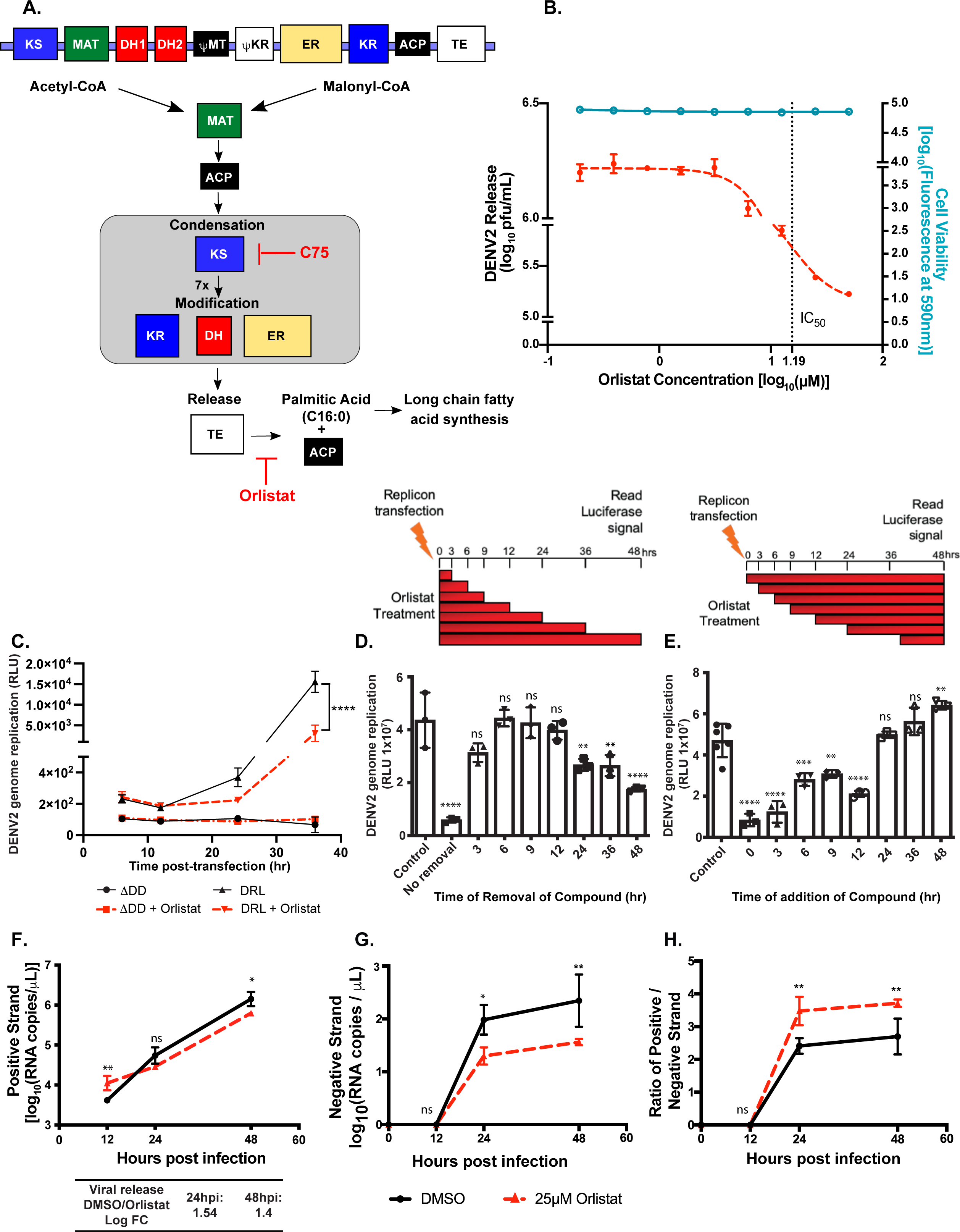
FAS activity is required for DENV2 RNA replication. (A) The domains of mammalian fatty acid synthase (FAS) and their role in enzymatic reactions with control points by C75 and Orlistat [adapted and modified from^11^]. KS, β-ketoacyl synthase; MAT, malonyl-CoA-/acetyl-CoA-ACP-transacylase; DH1 and DH2, dehydratase; ΨME, pseudo-methyltransferase, ΨKR, pseudo-ketoreductase; ER, β-enoyl reductase; KR, β-ketoacyl reductase; ACP, acyl carrier protein; TE, thioesterase. (B) Huh7 cells were infected with DENV2 16681 and treated with the indicated concentrations of Orlistat. Released virus was assayed by plaque assay and cytotoxicity was measured with resazurin. (C) Huh7 cells were electroporated with the indicated DENV2 luciferase replicons. After 2hr of incubation, media was changed and Orlistat or DMSO were added to the cells. Lysates were harvested at the indicated time points and assayed for luciferase expression. (D-E) Huh7 cells were electroporated with the WT DENV2 luciferase replicon. After 2hr of incubation, media was changed and Orlistat or DMSO were added to the cells. (D) Time of removal assay; Supernatant with Orlistat was removed at the indicated time points (post-transfection); cells were washed and media without the inhibitor was added to the cells. Lysates were harvested at 60hr post-transfection and assayed for luciferase expression. (E) Time of addition assay; After 2hr of incubation with the transfected replicon, media was changed and DMSO was added to the cells. At the indicated time points (post-transfection) the media was changed and media containing Orlistat was added. Lysates were harvested at 60hr post-transfection and assayed for luciferase expression. (F-H) Huh7 cells were infected with DENV2 and treated with 25μM Orlistat or DMSO. Supernatants and cell pellets were collected at the indicated time points. (Virus supernatants were titrated by plaque assay). (F) Intracellular DENV2 positive strand RNA was measured by qRT-PCR. (G) Intracellular DENV2 negative strand RNA was also measured by qRT-PCR. (H) The ratio of positive to negative strand DENV2 RNA at the indicated time points treated with Orlistat or DMSO. Two-way ANOVAs with multiple comparisons test were performed: *=p<0.05, **=p<0.01, ***p<0.001, ****p<0.0001; n=3 for all experiments.

Here, we demonstrate that Orlistat treatment impacts DENV2 viral RNA replication by disrupting negative-strand RNA synthesis. Orlistat also alters the virus-induced ER membrane structures both by reducing the number of vesicles in virus-infected cells and altering their morphology. Using metabolomics, we link these mechanistic changes in virus-infected cells to the chemical changes caused by Orlistat treatment. Finally, metabolic pathways do not act in a vacuum, but instead perturbation of one pathway can disturb other cellular pathways. One key pathway we discovered that was altered by Orlistat treatment of DENV2-infected cells was purine metabolism. We demonstrate that inhibition of purine metabolism in addition to FAS activity further disrupts viral replication. Essentially, we have identified the chemical composition of the metabolic environment that is refractory to viral replication whilst maintaining cellular viability. Understanding the metabolic effects of drug treatment on virus-infected cells is imperative to understand how metabolism can be exploited as an avenue for intervention.

## Results

### Early events in viral replication can be targeted by inhibition of FAS

To elucidate the role of FAS and its multiple intermediates and products during DENV2 replication, we took advantage of inhibitors of FAS activity. Both C75 and Orlistat reduced DENV2 release from Huh7 cells (^14, 16^ and Fig 1B). To further investigate the specific steps in the virus life cycle impacted by FAS inhibition we focused on Orlistat given its ability to block the final product of FAS. To distinguish the effect of Orlistat on viral RNA translation versus replication, we used two different DENV2 replicons. First, a DENV2 replicon with luciferase (DRL), which reports active protein translation and viral replication, and second a DENV2 replicon with luciferase and a defective polymerase (ΔDD). The ΔDD replicon, with it’s deletion in the active site (GDD) motif of the polymerase, is only capable of translating the transfected RNA and cannot replicate^144^. With DRL, we saw the translation of input RNA at 6hr with an increase in luciferase signal over time as the RNA is replicated (Fig 1C). However, in the presence of Orlistat, we observed reduced luciferase signal, indicating that DRL does not replicate as efficiently with less FAS activity. For ΔDD, at 6hr we observed translation of the input RNA, but (as expected) did not observe an increase in the luciferase signal over time due to the replication deficiency. When Orlistat was added to the replicon, we observed the same level of luciferase signal at 6hr, indicating Orlistat had no impact on translation of the input RNA. Therefore, inhibition of FAS reduced viral RNA replication but not translation.

The covalent modification of FAS by Orlistat is easily hydrolyzed due to the flexible hexanoyl tail of the compound^26^. Therefore, removal of Orlistat restores FAS activity and may restore viral replication. To test this hypothesis, we added Orlistat to the cells two hours after transfection of DRL. We removed the supernatant containing residual drug at the indicated time points post-transfection, then washed the cells and added new media without drug. We then measured luciferase signal from all samples after 60hr. Orlistat treatment for 48hr reduced DENV2 replication as expected (Fig 1D). Interestingly, virus replication was restored in samples where the drug was removed within 24hr. However, if the drug was removed at a later time point, viral replication was reduced compared to control. This result indicates that inhibition of FAS activity is most effective early following infection and that subsequent rounds of infection can be reduced by maintaining the compound for 24-48hr. To confirm this finding, we performed the complementary experiment where we electroporated the DENV2 replicon into Huh7 cells and overlaid the cells with normal media but replaced this with media containing Orlistat at the given time points post-transfection. Again, we observed that if the drug was added to the cells before 24hr, viral replication was reduced (Fig 1E). However, if we added the drug after 24hr, viral replication was not inhibited; further indicating that FAS activity was required early in infection but its modulation later in infection did not impact viral replication. Therefore, if FAS activity is restored early during the life cycle, DENV2 RNA replication can be restored. However, removal of drug at 36 or 48hr post-transfection seems to slow recovery. These data suggest that DENV2 RNA replication is directly dependent on the activity of FAS, and this dependency remains throughout the first 24hr of the life cycle.

### FAS inhibition disrupts DENV2 negative-strand RNA synthesis

Genome replication of positive-strand RNA viruses is dependent on making an uncapped negative-strand copy of the genome, which is used as a template to synthesize 5’ capped positive-strand genomes. Synthesis of the negative-strand occurs as a first step prior to synthesis of positive-strand genomes^27^. The ratio of positive to negative-strand viral RNA can serve as a marker for disruption of genome replication dynamics^28–30^. Thus, we tested the impact of Orlistat treatment on the levels of positive and negative-strand viral RNA over time to determine its impact on these processes. Huh7 cells were infected with DENV2 and treated at time 0 with Orlistat or vehicle control. Samples were collected at 0, 12, 24 and 48hr post-infection (hpi). Total RNA was extracted to measure the levels of positive and negative-strand viral RNA. We observed a minimal reduction in DENV2 positive strand RNA in vehicle versus Orlistat treatment (Fig 1F). However, we observed a significant reduction in viral negative-strand RNA compared to positive-strand RNA upon Orlistat treatment. This reduction was reflected in the ratio of positive to negative-strand RNA. The high levels of viral positive-strand RNA following Orlistat treatment could be due to its accumulation in the cell caused by a reduction in viral release (Fig 1B). These data indicate that FAS activity is important early during DENV2 replication for the production of the negative-strand replication intermediate.

### Inhibition of FAS does not impact virion stability and infectivity

We previously demonstrated that virion infectivity is linked to the lipid content of the ER^31^ and we demonstrated that inhibition of enzymes downstream of FAS that regulated unsaturated fatty acids (such as Stearoyl-CoA desaturase-1; SCD1) impacted particle infectivity, but inhibition of FAS by C75 did not significantly change infectivity^31^. Here we show that virus grown from Orlistat-treated cells also did not impact particle stability or infectivity.

We used thermal stability assays to determine if Orlistat impacted virion stability and infectivity. First, using equal PFU of virus grown with Orlistat or vehicle control, we maintained samples at 45°C for increasing periods of time and observed a similar loss in infectivity (Supplemental Fig 1A). We also subjected the virions to multiple rounds of freeze-thaw (at -80°C and at room temperature), and again saw no significant difference in infectivity of the two virus populations (Supplemental Fig 1B). Next, we tested the infectivity of virions generated in the presence and absence of Orlistat by quantifying their ability to re-infect fresh Huh7 cells. Using an equal MOI (as titrated on BHK-21 cells), we added both virion populations to fresh Huh7 cells. No significant difference in the titer of new viruses was observed (Supplemental Fig 1C). These data indicated that virions produced with Orlistat treatment had the same infectivity profile as control virions on mammalian cells. Similar results on virion stability were observed with C75 treatment (Supplemental Fig 1D and^31^). These observations further confirmed that the impact of FAS inhibition on DENV2 was at the RNA replication stage and not the virion assembly or maturation stage as previously observed for SCD1^31^.

### FAS Inhibitors disrupt DENV2-induced membrane architecture

One of the key questions in the field is whether the observed increase in FAS-mediated lipid biosynthesis has a direct link to the intracellular membrane rearrangements observed during DENV2-infection. To define this link, DENV2-infected and uninfected cells were treated with Orlistat, C75, or vehicle control and processed for thin-section electron microscopy (EM). Consistent with previous publications^7, 32, 33^ in DENV2-infected Huh7 cells, we observed a significant disruption of ER membranes and the formation of the well-known convoluted membrane (CM), vesicle (Ve), and vesicle packet (Vp) structures in the cytoplasm (Supplemental Fig 2). However, when cells were treated with FAS inhibitors, we observed significant differences in the phenotypes. When infected cells were treated with Orlistat, we observed structures with a similar diameter to virus particles aggregating in clusters in the cytoplasm but were not appear to be associated with or surrounded by cellular membranes (Fig 2A). Typically, virus particles in infected cells are seen enclosed in ER-derived membranes and traverse through the secretory pathway to exit the cell^7^. The CM, Vp, and Ve structures seen in infected cells (Supplemental Fig 2B) were not observed with Orlistat treatment (Fig 2A-C). We did, however, observe multiple large proteinaceous structures of unknown origin. These structures were also seen in uninfected cells treated with Orlistat (Fig 2D-F). With C75 treatment (Supplemental Fig 2C), we observed much more vacuolarization than normal uninfected Huh7 cells or untreated DENV2-infected cells (Supplemental Fig 2B). The phenotypes were also different to Orlistat-treated cells. Very few or no protein aggregates were observed in the C75-treated cells as opposed to the Orlistat treatment. CM similar to those observed in DENV2-infected cells were not observed. However, some possibly distorted Ve/Vp-like vesicles were identified (Supplemental Fig 2C). The extensive vacuolarization observed in infected cells treated with C75 was not observed in C75-treated uninfected cells (Supplemental Fig 2D).

**Fig 2.**
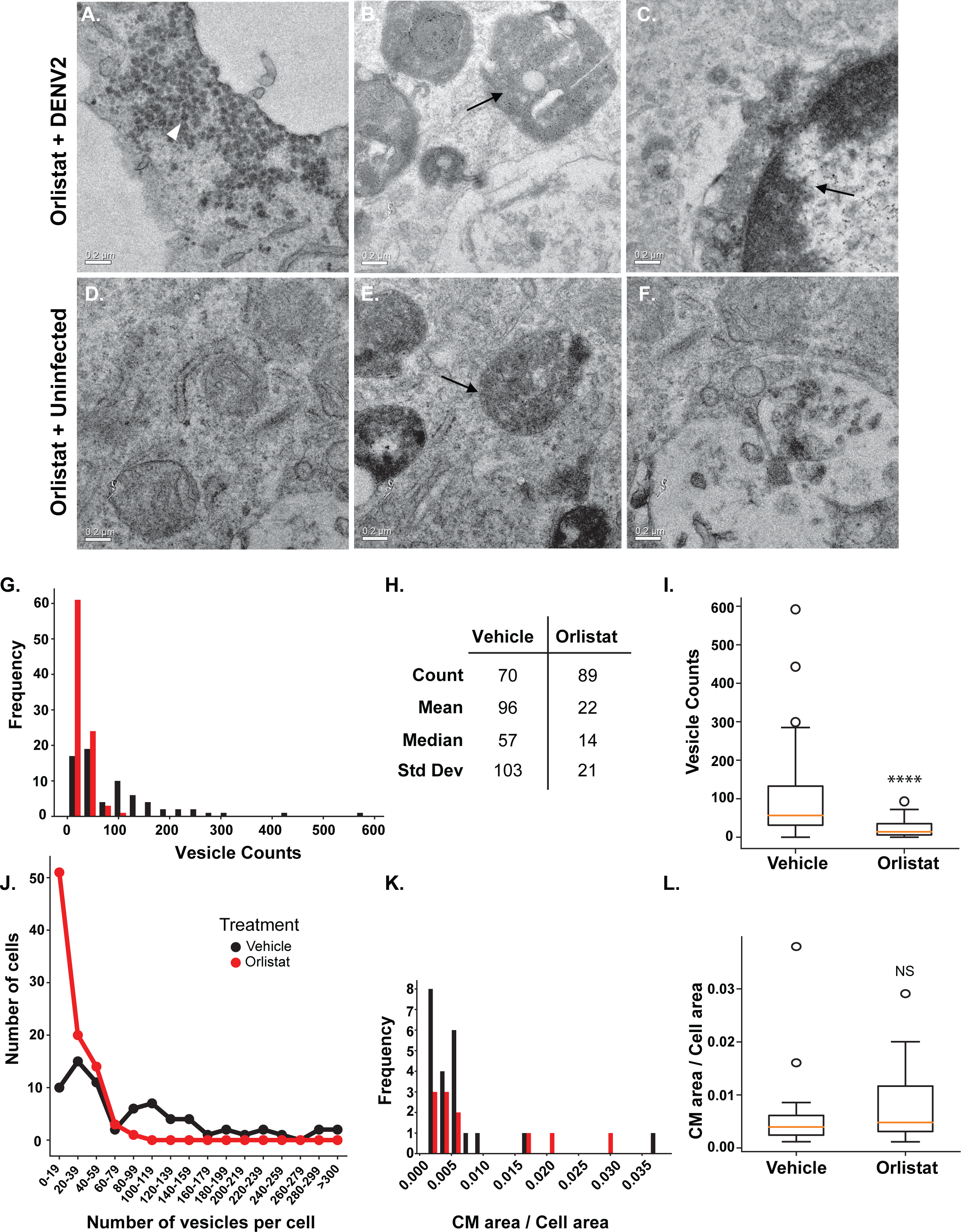
FAS inhibition disrupts DENV induced membranes. (A-L) Huh7 cells were infected with DENV2 (MOI=3) or mock infected and treated with Orlistat. At 24hr the cells were fixed and processed for EM imaging. All images are at 8800X magnification unless otherwise indicated. (A-C) DENV2-infected Huh7 cells treated with Orlistat (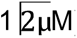). (A) White triangle marks virus like structures observed that are not associated with membranes. Membranes similar to those observed in DENV2 infected cells (ie: CM, Ve, Vp) are not observed with Orlistat treatment. (B-C) Many proteinaceous structures are present (marked with black arrows). (D-F) Uninfected Huh7 cells treated with Orlistat (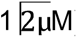). Similar proteinaceous structures as observed in B-C are also observed here (black arrow). (G-J) Montaged images of cell profiles were assessed for their vesicle content (G) A histogram of the number of vesicles in the montaged cell profiles with DENV2 infected Huh7 cells with Orlistat (shown in red) or vehicle treatment (black). (H) Summary of the numerical attributes of vesicle counts in each sample. (I) A box plot of vesicle count for the indicated samples displaying the median value with outliers denoted ****p=9.9e-12 from a Mann-WhitneyU non-parametric test (J) The distribution of vesicle counts in the same cells grouped by number of vesicles per cell. (K-L) The areas of convoluted membranes (CM) were measured in montaged images of cell profiles and compared to the area of the cell in the corresponding image. (K) A histogram of the relative area of CM in each sample. (L) A box plot of CM areas for the indicated samples displaying the median value with outliers denoted. NS: not significant, where p=0.22 from a MannWhitneyU non-parametric statistical test.

Since it is hypothesized that FAS-mediated lipid biosynthesis was elevated in DENV2-infected cells to provide substrates for virus-induced membrane structures^10, 14^, we expected the number of membrane structures to diminish upon FAS inhibition. To quantify the difference in treated versus untreated infected cells, we counted the virus-induced Ve and CM content in images of cell profiles (Fig 2)^34^. A histogram of the number of vesicles per cell profile demonstrates a non-normal distribution of the counts in both samples (Fig 2G). Infected cells treated with Orlistat have more cell profiles with fewer number of vesicles in them (Fig 2G). The median number of vesicles was significantly higher in vehicle-treated cells than Orlistat-treated (57 vs 14, respectively, Mann-Whitney U test, p < 0.001), showing a >75% reduction in vesicle number (Fig 2H, 2I). Large outliers in the DENV2-infected samples precluded use of parametric tests. The distribution of cells containing certain numbers of vesicles is shifted with Orlistat treatment. More cell sections had between 0-19 vesicles and no cell sections had more than 99 vesicles observed, whereas cell sections from the untreated cells had a wider distribution of vesicle counts and some cells with over 300 vesicles observed inside (Fig 2J). This indicates a drastic shift in the available replication sites per cell upon Orlistat treatment.

We also screened 40 cell sections per sample to examine the CM morphology and size. Morphologically, the CM appeared loosely packed and less distinctive in Orlistat-treated cells compared to vehicle control. The area of each CM was measured and normalized to the total cell area per section. The distribution of normalized CM areas in each sample was non-normal (Fig 2K), with no significant difference in the size of CM (Mann-Whitney U test, p = 0.22), (Fig 2L). Therefore, FAS inhibition by Orlistat results in distinct differences in the morphology of DENV2-induced CM structures but not in total area.

### Metabolomic profiling indicates significant chemical changes due to Orlistat treatment of DENV2-infected cells over time

Given the impact of FAS inhibition on the architecture of the virus-induced ER structures in DENV2-infected cells, we measured changes in the intracellular chemical composition to infer a direct link between lipid content and membrane structural alterations. We focused on Orlistat since it inhibits lipid biosynthesis through FAS as well as lipolysis through the inhibition of lipases. We infected Huh7 cells with DENV2 and treated them with Orlistat or vehicle control and collected the cells at either 6 or 18hpi (Fig 3A). The 6hpi time point represents very early events post-infection, and the 18hpi time point represents peak viral replication. Metabolites were extracted into polar and non-polar phases and high resolution liquid-chromatography-mass spectrometry (LC-MS) was performed in both positive and negative ionization modes for maximal coverage of the metabolome^9, 10^. The four data sets were analyzed to identify differences between Orlistat and vehicle-treated virus-infected cells at 6 and 18hpi. Molecular features (MF) (putative metabolites) observed were defined as unique mass to charge ratios (m/z) and retention times (rt). We generated linear models measuring the log fold change (logFC) in abundance of each MF. MFs were considered significantly different when the absolute log fold change was at least 1 and adjusted p-value was less than 0.005. A low p-value was used to decrease the likelihood of false discovery because of the large number of comparisons that were looked at simultaneously (Supplemental Table 1). Percent of features detected in the different phases and ionization modes are shown in Fig 3B.

**Fig 3.**
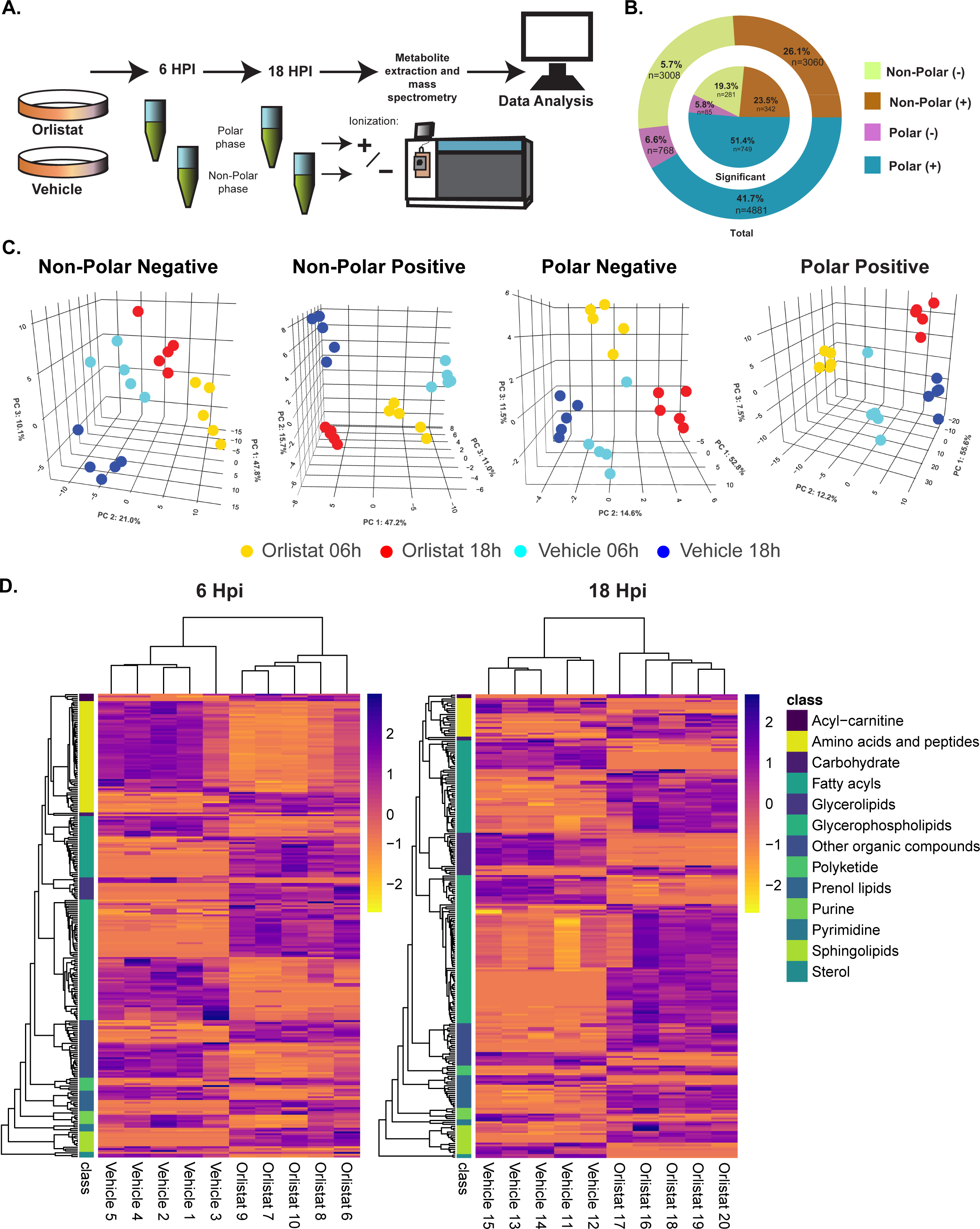
Orlistat treatment of DENV2-infected cells decreases key membrane lipid abundances. Huh7 cells were infected with DENV2 and treated with Orlistat or vehicle (n=5 for each). The cells were collected and processed for metabolomics at 6 and 18hpi. (A) An overview of the experimental procedure. (B) Pie charts of the total number of metabolites observed in each different mode after filtration steps (described in methods) (outer circle), and the number of metabolites found in each mode that were significantly different in at least one comparison after multivariate analysis (inner circle). (C) Principle component analyses (PCA) for the significantly different features found in each different mode. (D) Two-way hierarchical clustering of putatively identified metabolite abundances organized by lipid or metabolite class in Orlistat or vehicle treated samples at 6 and 18hpi.

We performed principal component analyses (PCA) on the significant MFs in all four phases and ionization modes. We observed a strong explanation (>50%) of the variation in the data due to time, regardless of treatment (Fig Supplemental 3). The main element that changes over time is the progression of viral infection. Hence, despite the drug treatment, the progression of infection changed the metabolic landscape of the cells. Additionally, we observed the most striking differences due to time and treatment in the non-polar metabolites with positive ionization, where principle components 1 and 2 explained 73% of the variation (Supplemental Fig 3). The change in non-polar metabolites in the positive ionization mode is expected given the mechanism of Orlistat, which targets fatty acid synthesis – a metabolic pathway that produces important precursors for many other non-polar lipids in the cell. However, we did see separation of all groups of samples by time and treatment when we also plotted the third principle component, indicating strong differences in the metabolic profile of the cells (Fig 3C).

### Orlistat treatment changes abundances of metabolite classes over time

The samples analyzed for metabolomics represent multiple different biological states. At the early time point post-infection, viral replication is low, therefore, the influence of infection on the metabolome is less and the influence of the drug is high. At the later time point, infection is attempting to ramp up but is modulated by Orlistat treatment. This dichotomy causes unique influences on the metabolome and highlights the impact of prolonged drug treatment. To characterize these different environments, we assigned putative annotations to the detected metabolites at MSI level 3 based on their accurate mass compared to the Human Metabolome Database (HMDB) or LIPID MAPS (Supplemental Table 1)^35–37^. We were able to assign putative identifications to 525 of the 1457 significantly altered metabolites (36%) to an accuracy of < 6 parts per million (ppm). Two-way hierarchical clustering of the putatively identified features showed a strong separation within lipid classes between treatment and time (Fig 3D).

#### Comparison of untreated samples to Orlistat-treated samples

Acyl-carnitines were all elevated with Orlistat treatment at both time points. A majority of the sphingolipids were also increased in abundance at both time points, with a minor fraction reduced in abundance post infection. A large number of amino acids and peptides showed reduced abundance in Orlistat-treated versus untreated samples at 6hpi. This distribution evened out at 18hpi. Fatty acids detected were also mixed in abundance, with some reducing in abundance and others increasing in abundance following treatment. The total number of fatty acyls significantly altered in abundance increased at 18hpi. A majority of the purines were also reduced at 6hpi with a minor fraction increased in abundance. At 18hpi, this ratio was more balanced. Prenol lipids and polyketides were also mixed in abundance trends at 6hpi with a majority increasing in abundance at 18hpi. Glycerolipids showed the most drastic response: all were elevated at the 6hpi time point, then showed a significant reduction at the 18hpi time point compared to untreated samples. In contrast, a majority of glycerophospholipids showed a significant reduction at the early time point compared to the later time point following treatment. Sterols were consistently reduced in abundance at both 6 and 18hpi post-treatment.

#### Comparison of Orlistat-treated samples at 6hr and 18hr post-treatment

We also compared the Orlistat-treated samples at the 6 and 18hpi time points to identify species that take longer to change in abundance and those that are quick to respond to treatment (Supplementary Fig 4). Compared to the 6hpi time point, treatment for 18hpi showed significant reduction in acyl-carnitines, amino acids/ peptides and a majority of the fatty acyls and glycerophospholipids. Addtionally, at 18hr, almost all glycerolipids also showed a drastic reduction with a very minor fraction showing an increase. This indicates that these molecules require a prolonged period of treatment to reduce in abundance. Prenol lipids showed more even distribution in abundance between the time points. A majority of purines, pyrimidines, sphingolipids and sterols were increased at 18hpi compared to 6hpi of treatment indicating that a reduction of these molecules occurs early following treatment and then recovers with time.

### Network analysis reveals changes to multiple biochemical pathways upon treatment of DENV2-infected cells with Orlistat

Metabolomic profiling provides data on altered chemical composition of cells and highlights a spectrum of changes across the cellular metabolic network. We performed pathway analysis to understand the diverse metabolic impacts of Orlistat treatment on cellular biochemical pathways during DENV2 infection. By comparing Orlistat to vehicle treatment at 6 or 18hpi, we identified multiple dysregulated biochemical pathways (Fig 4 and Supplemental Table 2). As expected, we observed a dysregulation in fatty acid biosynthesis and metabolism upon Orlistat treatment (Fig 4A). Interestingly, the most significantly dysregulated pathway at 18hpi with Orlistat treatment was purine metabolism (Fig 4A and Supplemental Table 2). The pathway analysis platform we utilized also assigns putative identifications (empirical features) to metabolites based on their relationships within a biochemical network (Supplemental Table 3)^38^. This platform complements our earlier analysis (Fig 3) as it identifies more metabolites than lipids. Visualizations of the biochemical networks show decreases in many empirical features at 6hpi in Orlistat vs vehicle treatments (Fig 4B), yet relatively few biochemical pathways were significantly disturbed at this time point (Fig 4A). At 18hpi the network displays more empirical features that are increased (Fig 4B), possibly to compensate for the reduction of *de novo* fatty acid synthesis.

**Fig 4.**
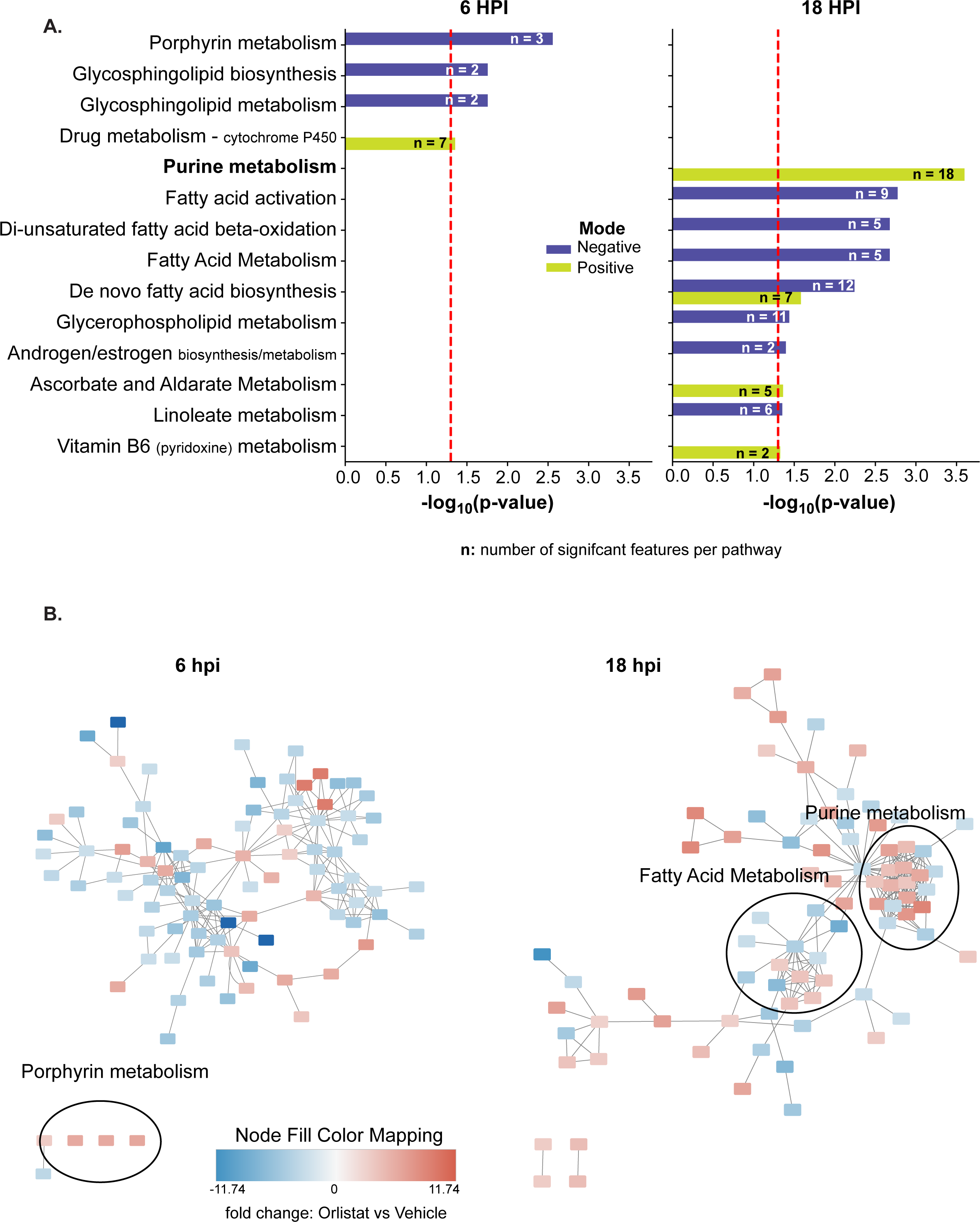
Network analysis reveals key metabolic pathway changes in Orlistat-treated DENV2 infected cells. Huh7 cells were infected with DENV2 and treated with Orlistat or vehicle. The cells were collected and processed for metabolomics at 6 and 18hpi. (A) Pathway analysis was performed on the features measured in negative and positive ionization modes from 6 and 18hpi. The indicated biochemical pathways were found to be significantly (p<0.05) disturbed at 6hpi and 18hpi in Orlistat versus vehicle treated DENV2-infected cells. Red dotted line is the threshold for significance at 0.05. (B) The networks of empirically identified chemical features at 6 and 18hpi. Features belonging to the most significantly disturbed biochemical pathways are indicated by ellipses. n: the number of features identified in each pathway.

### Orlistat-mediated metabolic disruptions in *de novo* fatty acid biosynthesis identifies purine metabolism as a key pathway for DENV2 replication

The purine metabolism pathway is responsible for maintaining the supply of adenine and guanine nucleotides in the cell. Nucleotide levels are controlled via a *de novo* pathway and a salvage pathway (Fig 5A). Changes in cellular purine levels likely have widespread impacts since they are key energy sources, components of DNA and RNA and cofactors for many cellular functions^39^. Our pathway analysis indicated an increase in *de novo* purine metabolism through the GMP branch as well as the salvage pathway (Fig 5A, B). We confirmed the identity (to MSI level 2) of select purine metabolites based on their LC-MS/MS spectra (Supplemental Fig 5)^36^. Specifically, we confirmed the identity of GMP, guanosine, AMP, adenosine and inosine. The abundance of these metabolites at 6 and 18hpi are plotted (Fig 5C). Our pathway analysis indicated a reduction in AMP and GMP with increases in adenosine, guanosine and inosine at 18hpi with Orlistat treatment (Fig 5C). This study does not address whether these changes are caused directly by Orlistat treatment or by the reduction in viral replication from Orlistat. To investigate this, we characterized the importance of the salvage pathway of purine metabolism during DENV2-infection. We used 6-thioguanine (6-TG), which is an FDA-approved inhibitor of purine biosynthesis used in the treatment of leukemia^40^. This drug competes with hypoxanthine and guanine for the activity of hypoxanthine-guanine phosphoribosyltransferase (HGPRT). Through inhibition of HGPRT, 6-TG reduces the salvage pathway of purine metabolism. Since we observed an increase in this pathway upon Orlistat treatment, we tested the impact of this drug on DENV2 replication. We observed a dose-dependent decrease in DENV2 replication with 6-TG treatment and almost no cellular cytotoxicity (Fig 5D). We then combined Orlistat and 6-TG, as well as C75 and 6-TG to treat DENV2-infected cells and interestingly found low cell cytotoxicity. However, in both cases we observed a greater decrease in viral replication than either one independently (Fig 5E). This indicated that DENV2 replication was further decreased when both fatty acid and purine biosynthesis were inhibited. Our study has demonstrated that utilizing one inhibitor to identify metabolically refractory environments can also lead to a discovery of another metabolic choke point for virus infection. Combining these two choke points can have greater impact on viral replication.

**Fig 5.**
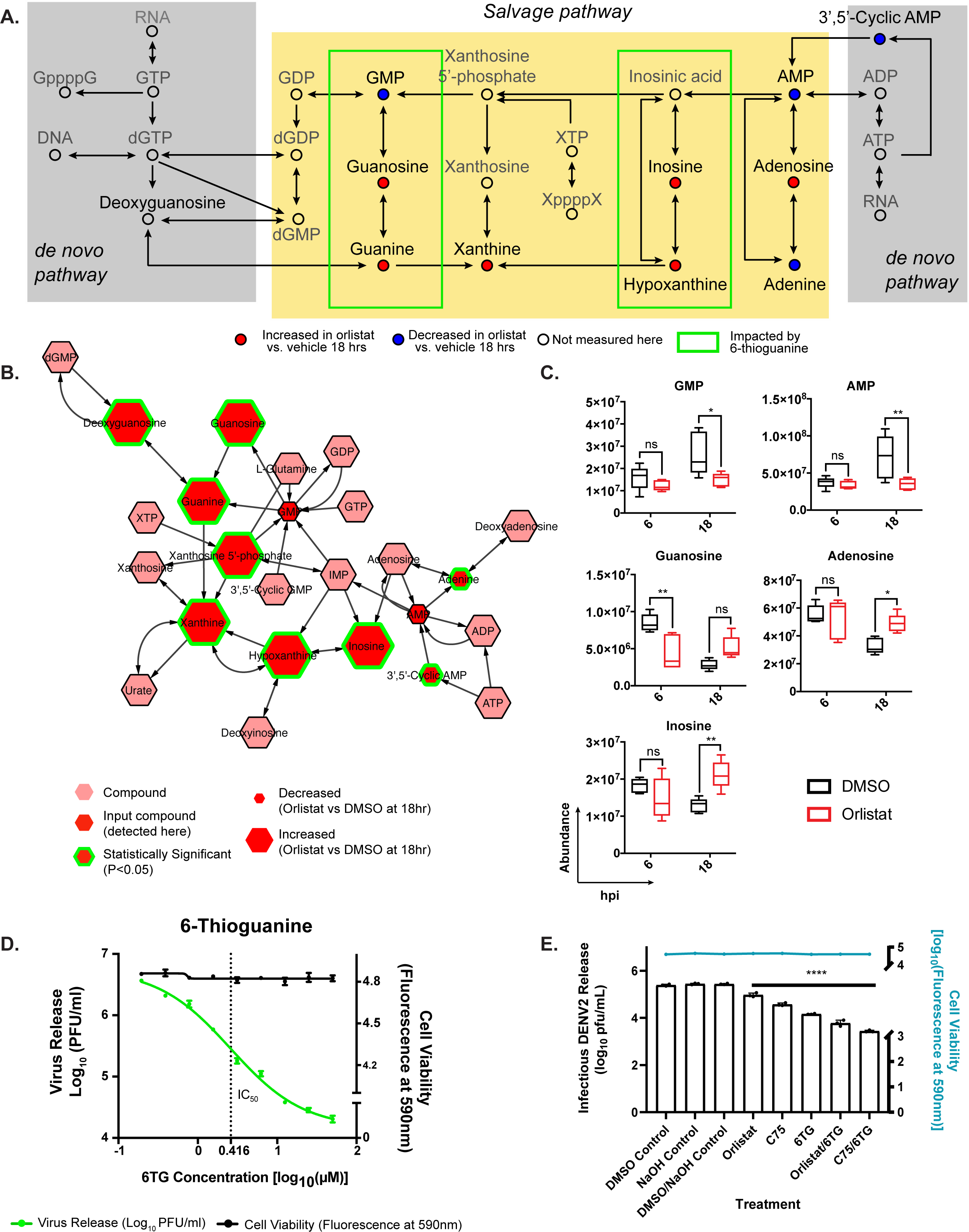
Orlistat treatment of DENV2 infected cells reveals a viral dependence on purine metabolism. (A) A map of purine metabolism based on the KEGG database^72^. Blue closed circles indicate metabolites that we putatively identified as decreased in Orlistat vs vehicle at 18hpi. Red closed circles indicate metabolites putatively identified as increased in Orlistat vs vehicle at 18hpi. Open circles are intermediate metabolites not detected here. The common name for the metabolite is listed above the circle. The green boxes are part of the pathway impacted by 6-TG treatment. (B) The purine metabolism network at 18hpi composed of metabolites significantly increased or decreased with Orlistat treatment as identified by Cytoscape. (C) The normalized abundance values (described in the methods section) for select metabolites in purine metabolism whose identifications were confirmed by MS/MS (shown in Supplemental Fig 5). Two-way ANOVAs with multiple comparisons test were performed. (D) Huh7 cells were infected with DENV2 or mock infected and treated with the indicated concentration of 6-TG. DENV2 release was measured after 24hr with plaque assays, and cytotoxicity was measured at 24hr with resazurin. Data was fit with a non-linear regression; n=3. (E) Huh7 cells were infected with DENV2 and treated with DMSO, NaOH, DMSO/NaOH, 15.5µM Orlistat, 2.6µM 6-TG, 3.7µM C75, 15.5µM Orlistat and 2.6µM 6-TG, or 3.75µM C75 and 2.6µM 6-TG. DENV2 release was measured at 24hr by plaque assays. A one-way ANOVA with a Dunnett’s multiple comparisons test was performed: *=p<0.05, **=p<0.01, ***p<0.001, ****p<0.0001; n=3. The stars indicate significance level in the multiple comparisons test for each sample compared to its corresponding vehicle control.

## Discussion

It has become increasingly clear that synthesis of cellular lipids is important for the replication of positive strand RNA viruses, but many questions remain surrounding the mechanisms and implications for viral replication. Here, using a clinically available inhibitor of FAS, we identifed a metabolic landscape in human cells that was refractory to viral infection but also identified compensatory pathways (such as purine metabolism) that allowed limited viral replication to occur. This led to identifying a combination of inhibitors that provided a synergistic reduction in viral replication. This work also demonstrated that there was a direct link between *de novo* fatty acid biosynthesis and the formation of viral-induced membrane structures, answering a long-standing question in the field.

Metabolomic profiling of DENV2-infected cells has been done previously by our group and others^9, 10, 41^. Here we present the first metabolomic analysis of DENV2-infected cells with an inhibitor targeting a key branch of cellular metabolism. This method allowed us to characterize the overall metabolic impact of FAS inhibition on DENV2-infected cells and identify additional biochemical pathways that were dysregulated. Purine metabolism was the most dysregulated pathway upon Orlistat treatment of DENV2-infected cells. In particular, we found an increase in *de novo* purine metabolism through the GMP branch as well as the salvage pathway.

There are likely many causes for dysregulated purine metabolism with FAS inhibition. Here, we will discuss two potential models: (i) a feedback loop that activates acetyl CoA-carboxylase (ACC) and (ii) an accumulation of acetyl-CoA that is shunted towards the TCA cycle. **Model (i):** First, when FAS activity is inhibited, guanine levels may be increased in order to regulate ACC activity (the rate-limiting enzyme upstream of FAS) in an effort to force more fatty acid synthesis. Guanine nucleotides increase ACC activity *in vitro*, hence they could increase ACC activity in the cell^42^. We and others have previously shown the importance of ACC activity during flavivirus infection^12^. C75 treatment causes an initial activation and futile cycling of malonyl-CoA production followed by a decrease in ACC activity. This was shown to result in an initial decrease in ATP followed by an activation of AMPK and the production of ATP through catabolic reactions^43^. Hence, inhibition of FAS via Orlistat could cause a similar dysregulation of ACC activity and the initial decrease in ATP as we see here. However, others have demonstrated that Orlistat treatment reduces AMPK activation due to its inhibition of lipolysis^44^. Perhaps the dual action of Orlistat to inhibit FAS resulting in a disturbance of ATP levels coupled to the inhibition of lipolysis and reduction of AMPK acts to further suppress ATP levels. The increase in guanine may represent an effort of the virus or cell to increase ACC activity or simply to activate the purine metabolism salvage pathway in order to increase purine nucleotide levels in the cell. The activity of HGPRT during viral infection and/or Orlistat treatment should be further characterized to shed light on this hypothesis.

### Model (ii)

Another consequence of FAS inhibition is the accumulation of acetyl-CoA that is not converted into fatty acids. This pool of acetyl-CoA could alternatively be redirected into the TCA cycle. During the TCA cycle, either ATP or GTP may be produced by succinyl-CoA synthetase (SCS). The liver is a major site of nucleotide synthesis^45^, and the human liver has more GDP-specific SCS than ADP-specific SCS^46^. This would favor the production of GTP and the subsequent increase in deoxyguanosine at 18hpi allowing for the formation of the other guanine-based compounds. This could also explain the decrease in adenine-based compounds at 18hr. The guanine-based and adenine-based compounds can cycle through the salvage and *de novo* purine synthesis pathways to generate more nucleotides for virus replication.

We observed that FAS activity is particularly important for DENV2 replication at early timepoints of infection when RNA replication complexes are forming and producing negative strands of RNA. The need for early lipid synthesis is supported by our work on SCD1, another lipogenic enzyme^31^. A similar need for fatty acids early in viral replication was observed in Rift Valley fever virus infection where cells restrict viral replication by increasing AMPK activity to reduce fatty acid synthesis^47^. The addition of exogenous fatty acids rescued viral replication despite the cellular efforts; however, this only worked if palmitic acid (C16:0) was added at an early time point or before viral replication began^47^. Early synthesis of fatty acids could be common theme for RNA viruses that need to ensure an adequate concentration of required substrates for the construction of membranous replication complexes.

The relocation of FAS to the viral replication complex suggests a need for fatty acid substrates at the site of viral replication^14^. However, other lipid composition requirements of the replication complex and the impact of the lipid composition on viral replication dynamics is unknown. The data presented here suggests that when fatty acid synthesis is reduced, viral RNA replication dynamics are slowed. Orlistat treatment resulted in a slight reduction of positive strand RNA synthesis, but negative strand synthesis was severely reduced. This is consistent with slowed early replication under Orlistat treatment and suggests that negative strand synthesis has a unique dependence on membrane fatty acid composition and/or concentration from FAS activity. NS5 is the RNA dependent RNA polymerase that synthesizes negative strand viral RNA early during viral infection and later switches to positive strand synthesis. These observations could suggest that early NS5 activity is dependent on fatty acid concentration or FAS activity. Then, once the synthesis of negative strands is established, RNA replication can proceed irrespective of FAS activity. This dependence on FAS activity may be due to a total reduction in available membranes to establish replication sites, or it may be due to a reduction in complex lipid species that require fatty acids and can generate specific characteristics in the membranes.

Importantly, we observed distinct changes to the hallmark membrane rearrangements that occur in DENV2-infected cells upon inhibition of FAS. The pattern of changes to the membranes differed between Orlistat and C75 suggesting that the outcomes correlated with the mechanism of action of each compound. The different phenotypes observed may indicate that differential inhibition of FAS (at different subdomains of the enzyme) impacts the formation of these membranes directly or may represent off-target metabolic effects of these compounds on membrane development. In addition to inhibition of FAS, Orlistat is a lipolysis inhibitor that binds to lipases preventing the hydrolysis of triglycerides^22^. This alternate function of Orlistat may help drive the observed impact on membrane formation. The importance of fatty acid synthesis in viral-induced membrane rearrangements has also been observed with inhibition of acetyl-CoA carboxylase (ACC) during West Nile virus infection. Inhibition of ACC disrupted the formation of membranous complexes in virus-infected cells as seen in EM images^12^. ACC catalyzes the carboxylation of acetyl-CoA to produce malonyl-CoA, which is a substrate of FAS, hence inhibition of ACC also reduces fatty acid substrates in the cell. Together, these data demonstrate that lipid composition and *de novo* synthesized fatty acid substrates are critical for the formation of viral-induced specialized membranes in the ER.

Our findings presented here provide a direct link between DENV2-induced cellular metabolic changes and ER membrane rearrangements that are dependent on FAS activity. These membranes are critical as platforms for viral replication and to protect the replicative intermediate from the cellular immune system. We further demonstrated that a change in FAS activity impacted viral RNA replication dynamics and we propose that this is due to changes in lipid composition of the ER membranes that are critical to house the viral replication machinery and may impact the viral polymerase activity. Finally, we demonstrated that inhibition of FAS activity in DENV2-infected cells resulted in dysregulation of purine metabolism. This finding highlighted the importance of HGPRT in DENV2 replication independent of FAS activity. Hence, we have confirmed the potential to re-purpose Orlistat as an anti-viral, demonstrated its mechanistic impact on viral replication dynamics and identified HGPRT as another antiviral target. Furthermore, mechanistic insight into the relationship between FAS and purine metabolism will likely identify further antiviral targets and illuminate the interconnectedness of cellular metabolic pathways.

## Materials and Methods

### Cell lines, viruses and inhibitors

The cell lines used for this study were as follows: BHK-21, Clone 15 (ATCC CCL-10); and Huh7 (From Dr. Charles Rice,^48^). Huh7 cells were maintained in Dulbeccos Modified Eagle Medium (DMEM) (Gibco, LifeTech), while BHK-21 were maintained in Minimum Essential Media (MEM) (Gibco, LifeTech), both were supplemented with 0.1 mM nonessential amino acids, 0.1 mM L-glutamine, and 10% Fetal Bovine Serum (Atlas Biologicals) at 37°C with 5% CO_2_.

DENV2 (16681)^49^ passaged in C6/36 cells was used for the experiments described here. Infectious virus titers were quantified by plaque assay on BHK-21 cells as described previously^50^. The DENV2 replicon used for this study was described previously^14^, where the structural proteins of DENV were replaced with a *Renilla luciferase*.

The inhibitors used for this study were as follows: Orlistat (Sigma-Aldrich) freshly diluted in DMSO; C75 (Sigma-Aldrich) diluted in DMSO and 6-Thioguanine (6-TG) (Sigma-Aldrich) diluted in 1M NaOH. All inhibitors were further diluted to the indicated concentrations in DMEM before being added to cells. Solvent concentrations were maintained at or below 0.1%. Huh7 cells were infected with virus as described above and overlaid with the indicated concentrations diluted in DMEM. Supernatants were collected at 24hpi and plaque assays performed. Cytotoxicity was measured by adding resazurin [alamar blue (ThermoFisher)] diluted to 1x in DMEM according to the manufacturor’s protocol and incubated on cells for 2-4hr and reading fluorescence on a Victor 1420 Multilabel plate reader (Perkin Elmer) with excitation at 560 nM and emission at 590 nM.

### RNA extraction and qRT-PCR

RNA was extracted from cells with Trizol (ThermoFisher) and from virus in supernatant using Trizol LS (ThermoFisher). A one-step qRT-PCR kit with SYBR green from Agilent was used. Reactions were set up according to the manufacturer’s protocol and run on a LightCycler 96 real-time PCR machine (Roche). The cycling parameters were: 20 mins at 50°C for reverse transcription, 5 min at 95°C, 45 two-step cycles of 95°C for 5 seconds and 60°C for 60 seconds, and a melt curve starting at 65°C and ending at 97°C. DENV2 primers^51^ were used to quantify viral RNA copies by comparing to a standard curve of *in vitro* transcribed viral RNA from a DENV2 cDNA subclone^52^. The DENV2 negative strand RNA was measured with a two-step qRT-PCR described previously ^53^.

### Primers

DENV2 Positive strand:

Forward: ACAAGTCGAACAACCTGGTCCAT Reverse: GCCGCACCATTGGTCTTCTC

DENV2 Negative strand:

Forward: CGGTCATGGTGGCGAATAA Reverse: CATTCCATTTTCTGGCGTTCT

### Measurement of virus infectivity and stability

Virion infectivity was measured via plaque assays described previously^31^. Briefly, the supernatants from inhibitor-treated cells were collected and titrated on BHK-21 cells. Using an equal MOI, we added both virion populations to new Huh7 cells and allowed the virus to adsorb for one hour. Given the potential for unmetabolized Orlistat in the virus sample, we added the original concentration of Orlistat to the vehicle control sample during the attachment stage. We then washed the unattached virus, overlaid the cells with new media without drugs, allowed the virus to replicate for 48, and titrated the new supernatant on BHK-21 cells. For the thermal stability assays, the supernatants from inhibitor-treated cells were collected, titrated and diluted to the indicated concentration. The virus samples were incubated at the indicated temperatures for 15 minutes, re-equilibrated to room temperature, and immediately titrated. Alternatively, the virus samples were subjected to rounds of freezing at -80C or thawing at RT and titrated on BHK-21 cells after each round.

### Cell sectioning for Electron Microscopy

Huh7 cells were infected with DENV2 at MOI = 3 and treated with the indicated compounds or vehicle control. At 24hpi, supernatant was collected for plaque assay and cells were fixed in a solution containing 2% glutaraldehyde in cacodylate buffer (0.1M Na-cacodylate, 2mM MgCl_2_, 2mM CaCl_2_, 0.5% NaCl, pH 7.4) for 10 minutes, rinsed twice in cacodylate buffer followed by post-fixation for 10 minutes in a reduced osmium solution (1% OsO_4_ + 1.5% potassium ferrocyanide). Osmium exposed samples were rinsed 3 X with distilled water and scraped from the plate into 500 μl volume. One ml of 1.5% low-melt agarose held at ∼45°C was gently mixed with the cell volumes and spun down at 3300xg in a microcentrifuge for 10 min at room temperature. Agarose was then cooled in the freezer for 10 minutes to allow it to solidify. Cell pellets were cut from the agarose blocks and placed into scintillation vials for dehydration in a graded ethanol series. Each step of the series was allowed to diffuse into the pellets for 20 min/step; steps consisted of 10%, 30%, 50%, 70%, 90%, 100% ethanol in water followed by two 20 min steps in pure acetone. Following dehydration, cell pellets were infiltrated in a graded durcupan-epoxy resin series. (Durcupan-epoxy recipe: 2.0 g Epoxy Embedding medium, 2.6 g Component A, 4.9 g Component B, 0.2 g Component C, and 0.3 g Component D). The infiltration steps used were: 25% durcupan-epoxy:acetone-2h, 50%-2h, 75%-overnight (approx. 16h), and 100% for 6 hours. Post-infiltration, pellets were put into block molds with labels and polymerized in a 60°C oven for 72 h. After 72 h blocks were removed from the oven and 90 nm sections were cut on a Leica UC7 Ultramicrotome and collected onto formvar-carbon coated copper slot grids. These grids were post-embed stained with 2% uranyl acetate for 5 min and Sato’s lead for 1 min.

Grids were initially evaluated on the CM-200 transmission electron microscope followed by montage collection of cell profiles using SerialEM on an FEI Technai T20 200kV electron microscope. Sixty-nine cell profiles were collected of the DENV2 infected Huh7 cells while 88 cell profiles were collected of DENV2+Orlistat exposed cells. Image acquisition occurred at a nominal magnification of 5000X on a 2K CCD camera from Gatan. Montaged images were viewed using the IMOD software package. Vesicles were counted manually on a per cell profile basis. CM area was measured by manually outlining the area and computing the values using IMOD. Descriptive statistics and hypothesis testing was performed with NumPy and SciPy in Python^54^.

### Metabolite extraction

For the metabolomics experiments, we used five biological replicates for each time point and treatment. Cells (1.5 x10^6^) were harvested at the indicated time point and metabolites were extracted from an equal number of cells per sample with a modified Bligh and Dyer protocol^55^. Briefly, a mixture of 2:1 chloroform methanol, 0.1% acetic acid and 0.01% butylated hydroxy toluene (BHT) was added to the cell suspension in ammonium bicarbonate to generate a 4:1 ratio of organic solvent to cells. Following lipid extraction, the organic phase was separated from the aqueous phase by centrifugation and each were dried down under a N_2_ stream in low retention microfuge tubes (Fisher). The dried lipids were resuspended in 75 μl of ice-cold methanol and vortexed for 10 s.

The samples were then centrifuged at 13,400×g for 5 min to remove any particulates. To prevent degradation of metabolites, samples were processed on ice or at 4°C throughout the process. Samples from aqueous and organic phases were dried separately by speed-vacuum centrifugation and were stored at -80°C until metabolomics analysis. Prior to LC-MS analysis, aqueous and organic extracts were resuspended in 75 µL of 50% methanol and methanol, respectively. Resuspended samples were centrifuged and transferred into LC-MS vials and ready for analysis.

### LC/MS analysis

One technical replicate from each sample was analyzed on a LTQ Orbitrap XL instrument (Thermo Scientific, Waltham, MA). It was coupled to an Agilent 1100 series LC (Agilent Technologies, Santa Clara, CA) equipped with a refrigerated well plate auto sampler and binary pumping device. Reverse-phase liquid chromatography was used to analyze the samples in both phases.

#### Polar metabolite analysis

An Atlantis T3 column (Waters Corp., Milford, MA 1.0 x 150 mm, 5.0 μm) was used for the LC separation. Solvent A was water + 0.1% formic acid. Solvent B was acetonitrile + 0.1% formic acid. The flow rate was set to 140 μL/minute. A sample volume of 5 μL was loaded onto the column. The gradient was as follows: time 0 minutes, 0% B; time 1 minutes, 0% B; time 41 minutes, 95% B; time 46 minutes, 95% B; time 50 minutes, 0% B; time 60 minutes 0% B. We ran the LC-MS analysis twice, using positive and negative polarity electrospray ionization (ESI). Data were acquired using data dependent scanning mode. FTMS resolution of 60,000 with a mass range of 50– 1100 was used for full scan analysis.

#### Non-polar metabolite analysis

An Xterra C18 column (Waters Corp., Milford, MA, 2.1 x 150 mm, 5.0 μm) was used for the separation of the non-polar metabolites. Solvent A consisted of water + 10mM ammonium acetate + 0.1% formic acid. Solvent B was acetonitrile/isopropyl alcohol (50/ 50 v/v) + 10mM ammonium acetate + 0.1% formic acid. The flow rate was 300 μL /minute. A sample volume of 10 μL was loaded onto the column. The gradient was as follows: time 0 minutes, 35% B; time 10 minutes, 80% B; time 20 minutes, 100% B; time 32 minutes, 100% B; time 35 minutes, 35% B; time 40 minutes 35% B. The LC-MS analysis was run twice, with both positive and negative polarity ESI. The acquired data were evaluated with Thermo XCalibur software (version 2.1.0).

#### Metabolite Identification

For identification of select metabolites, accurate mass (*m/z*) and elution profile of metabolites in reverse-phase liquid chromatography were taken in account. MS/MS spectra of metabolites generated in data dependent scanning were manually interpretated and compared with publicly available databases (Human Metabolome Database, Metlin^56^) for identification of statistically significant molecules as described in the sections below.

### MS data processing and analysis

We converted raw mass spectrometry data to mzXML format with msConvert^57, 58^. Preprocessing and analysis were conducted in R open software. Peak picking was accomplished with the XCMS package using the centWave^59–61^ algorithm with a Gaussian fit for peak-picking and the OBI-Warp method for retention time correction and alignment^62^. Parameters used for XCMS were optimized using the IPO package^63^. Intensities for peaks were determined and normalized using the median fold change method^64, 65^. Features were removed if their retention time was outside of acceptable limits: 2–34 minutes for nonpolar modes and 2–48 minutes for polar modes. Isotopes were identified using the CAMERA package in R, and removed. We assumed values for features with no detectable peak in more than half of the samples in a group to be below the lower limit of detection and imputed with one-half the overall minimum intensity value. Groups with values for at least half of the samples in that group were imputed with the fillPeaks function of XCMS and normalized with previously calculated normalizing constants^66^. Where fillPeaks resulted in a zero intensity, we assumed the intensity is below the lower limit of detection, and imputed values with one-half the overall minimum value. These abundance values were used for pathway analysis described below.

Each of the four chemical and analytical modes (polar/non-polar and negative/positive mode) were processed and analyzed separately. To test for univariate dysregulation of metabolites among the 4 groups in each mode, we used the limma package in R^67^. This approach fit a linear model to each feature and makes adjustments in the t-statistic using an empirical Bayes approach. P-values were adjusted for false discovery rate in each pairwise comparison separately. A *p*-value of 0.005 was used as a cutoff because of the large number of comparisons that are looked at simultaneously in this study.

Heat maps were plotted with the pheatmap package in R, using an average linkage for hierarchical clustering, with each lipid group clustered individually, and peak areas scaled row-wise ^68^.

Molecular features (MFs) were identified as described previously^9^. Briefly, mass to charge ratio (m/z) were searched against the Human Metabolome Database (HMDB) and LIPID Metabolites and Pathways Strategy (LIPID MAPS). [M+H]+, [M+NH4]+ and [M+Na]+ adducts were accounted for calculating the neutral mass in the positive ionization mode and [M-H]- was accounted for that of the negative ionization mode. Only MFs with mass accuracy < 6 ppm error were further classified into metabolic classes. Lipids were classified according to the lipid classification system from LIPID MAPS^69^.

### Pathway analysis

We used a Student’s t-test on the imputed abundances for all the features at each time point with each treatment generating two comparisons of features in negative mode and two comparisons of features in the positive mode. Network analysis and metabolic pathway analysis was performed using the *mummichog* software version 2.0^38^. This platform tests the enrichment of input features against random data resampled from the reference list and produces an empirical *P* value for each pathway. Input metabolites were then annotated as empirical compounds. Metabolites with significant changes (p<0.05) were mapped onto metabolic pathways, linked in a network fig by known metabolic reactions and visualized in Cytoscape version 3.7.1^70^ (Fig 4B) or MetScape version 3.1.3^71^ (Fig 5B).

### Statistical Methods

The details of the statistical tests used here are noted in the figs and/ or in the corresponding fig legends. Briefly, drug inhibition studies (Fig 1 and 5) were analyzed with a non-linear regression using GraphPad Prism version 7.00 for Mac OS x (GraphPad Software, La Jolla California USA) to calculate the IC50. Differences in luciferase signal from the replicons (Fig 1) was calculated with a one- or two-way Analysis of Variance (ANOVA) with a Bonferroni multiple-comparison test in the GraphPad Prism version 7.00. Reduction in negative- and positive-strand RNA (Fig 1) was tested using two-way ANOVA with a Dunnett’s multiple-comparison test in GraphPad Prism version 7.00. Descriptive statistics of the vesicle counts and CM area from thin-layer microscopy images (Fig 2) was performed with NumPy and Python^54^. While hypothesis testing was accomplished via a non-parametric Mann-WhitneyU using SciPy in Python^54^. Statistical analysis for metabolomic data was as described in the sections above. Statistical comparisons of confirmed metabolites in purine metabolism (Fig 5) was accomplished using a two-way ANOVA with a Dunnett’s multiple-comparison test in GraphPad Prism version 7.00. And the inhibitor combination experiment (Fig 5) was analyzed with a one-way ANOVA with a Dunnett’s multiple-comparison test in GraphPad Prism version 9.2. Non-linear regression with four parameters was performed on the thermorstability data set of DMSO vs Orlistat treated virus (Supplemental Fig 1). The 95% confidence intervals for the LogIC50 values overlapped for each data set indicating that they are not significantly different. A linear regression of the freeze-thaw experiment (Supplemental Fig 1) was perormed in GraphPad Prism version 7.00. Each study shown is representative of at least two independent experiments.

## Supporting information

Gullberg et al_Supplemental Figures

Gullberg et al_Supplementary Table 1

Gullberg et al_Supplementary Table 2

Gullberg et al_Supplementary Table 3

---

**Supplemental Fig 1. DENV2 virion particle infectivity is not impacted by Orlistat treatment.** (A) Huh7 cells were infected with DENV2 and treated with Orlistat or DMSO. Virus supernatant was collected at 24hr and titrated by plaque assay. Each sample was diluted to equal titers incubated at 45°C for the indicated times and titrated on BHK-21 cells to measure the thermostability of the infectious particles. The results were fit with a four-parameter non-linear regression. (B) Virus samples were subjected to freeze-thaw cycles. The infectivity of the virus particles following each cycle was quantified. A linear regression showed that each slope does not significantly deviate from zero (DMSO p = 0.0621; Orlistat p = 0.3064). (C) Equal PFU of released DENV2 following treatment with Orlistat or DMSO was used to infect new Huh7 cells. After attachment of the virus the inoculum was washed, and the cells were overlaid with media without the inhibitor or DMSO. After 48hr the supernatants were collected and titrated. A student’s t-test was performed: ns; not significant: p>0.05. (D) A similar experiment to A was carried out with C75 and DMSO control. Each sample following C75 or DMSO treatment was diluted to equal titers, held at 45°C for the indicated times and titrated to measure the thermostability of the infectious particles. n=3 for all experiments.

**Supplemental Fig 2. DENV2 infected Huh7 cells with or without C75 treatment display normal rearrangements of cellular structures.** Huh7 cells with the indicated treatments were fixed and processed for EM imaging. All images are at 8800X magnification. (A) Uninfected Huh7 cells. (B) Huh7 cells were infected with DENV at MOI = 3. Samples were prepared for thin section EM at 24hr post-infection. Typical DENV- induced membranes are indicated^32^. CM; convoluted membranes, Vp; vesicle packets, Ve; vesicles within vesicle packets, Vi; virus particles. (C) DENV2 infected Huh7 cells treated with C75 (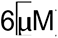). (D) Uninfected Huh7 cells treated with C75 (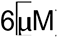). (E) Montaged images of cell profiles were assessed for their vesicle content Vesicles and convoluted membranes in montaged cell profiles of DENV2-infected Huh7 cells +/- C75 treatment.

**Supplemental Fig 3. Principle component analyses (PCA) shows segregation of the global metabolite profiles.** PCA plots for each phase and ionization mode. The variables compared in each graph are labeled. Left panels show differences in treatment, while right panels show time and treatment. Ovals around the data represents the 95% confidence interval of those samples (open circles or triangles) around the mean (solid circle). DENV2; dengue virus serotype 2, O; Orlistat, V; vehicle (DMSO), 06; 6hpi, 18; 18hpi.

**Supplemental Fig 4.** Two-way hierarchical clustering of putatively identified metabolite abundances organized by lipid class in Orlistat samples comparing 6 to 18hrs.

**Supplemental Fig 5. Identification of purine metabolites.** LC-MS extracted ion chromatograms, MS spectra and MS/MS spectra along with collision-induced fragmentation patterns of (A) adenosine monophosphate, (B) adenosine, (C) inosine, (D) guanosine monophosphate and (E) guanosine.

**Supplemental Table 1.**

**Metabolites detected in DENV2 infected cells with differential levels over time or upon treatment with Orlistat or Vehicle.** Metabolite abundance was obtained after features were filtered through our metabolomics pipeline as described in methods section. Features that displayed significantly different levels over time or upon treatment conditions were retained. Tab 1 of the table are features that were assigned putative identifications (MSI level 1-3) while tab 2 are features not reliably assigned an identification (MSI level 4). The following information is provided for each feature: M/Z_Avg_: the average mass to charge ratio for the five replicate samples. RT_Avg_: the average retention time of the five replicates in seconds. Detection mode: including the polar or non-polar phase of the sample and positive or negative ionization mode on the instrument. Name: the identification of the feature. Lipid class: The class of lipid according LIPID maps classification^37^. Formula: the chemical constituents of the feature at their neutral mass. Adducts: Any molecular protonated or deprotonated ions: [M+H]+, [M+NH4]+, [M+Na]+, [M-H]-. PPM error: the parts per million difference between the experimental mass and expected mass. Match ID: the accession number for each metabolite coming from the LIPIDMAPS database or Human Metabolome Database (HMDB). Number of alternate IDs: the number of other metabolite names that are isomers of the chosen name for the chemical formula. Log_2_ fold change: the Log_2_ fold change in abundance between 6 or 18hpi or between Orlistat treatment compared to Vehicle. Adjusted p-value: the adjusted P-value for the indicated comparison- described in methods.

**Supplemental Table 2. Dysregulated biochemical pathways in DENV2 infected cells following Orlistat treatment as determined through Mummichog pathway and network analysis of metabolomics data.** All features identified in the metabolomic dataset were analyzed with the mummichog platform^38^ and the identified pathways are provided here. The following information is provided for each pathway: Pathway: the name of the biochemical pathway. P-value- the p-value indicating the significance changes in the pathway above background levels. Features (KEGG id): the identification of the features according to the KEGG database that are found in the indicated pathway. Features (Name): The name of the corresponding features found in the pathway. Mode: the negative or positive ionization mode of the features identified in the pathway. Time: the hours post infection where the features were found different between treatment groups.

**Supplemental Table 3. Empirical metabolic features in DENV2 infected cells following Orlistat treatment as determined through mummichog pathway and network analysis of metabolomics data.** The features in our metabolomic dataset were subjected to network analysis with the mummichog platform, which assigns the most likely identification of the feature based on its location within a network. The following information is provided for each feature: Time: the hours post infection where the feature was found with different levels between treatment groups. Mode: the negative or positive ionization mode where the feature was measured. Empirical Compound: the internal identification according to the mummichog platform. KEGG ID: the accession number of the features according to the KEGG database. Name: the identification of the feature. Other IDs: the name of other metabolite that are isomers of the chosen name for the chemical formula. m/z: the average mass to charge ratio over the five biological replicates of the indicated feature. Retention time: the average retention time in seconds over the five biological replicates for the feature. Adduct: Any molecular protonated or deprotonated ions: M+H[1+], M[1+], M(C13)+H[1+], M+Na[1+], M+Na[1+], M+HCOONa[1+], M+Na[1+], M+H[1+], M+HCOONa[1+], M+Na[1+], M+2H[2+], M[1+], M(C13)+2H[2+], M+HCOONa[1+], M+H+Na[2+]. Mz difference: the difference between the experimental mass and expected mass. Statistic: the t-score calculated for each feature between orlistat treatment and vehicle. Significant: determined based on the p-value from the Studen’t t test.

## Author Contributions

RCG, NC, LSC, EL, THE, SK, CT, KK and RP carried out the experiments. MNI and AHJ carried out the mass spectrometry analyses. BG carried out the biostatistics. RCG, NC, LSC, EL, THE, SK, CT, KK, MNI, AHJ and RP wrote the manuscript. DJL, JT, RJK and RP helped with conceptualization and critical evaluation of data and critical reading of the manuscript.

