## Supplementary material for "Shifting a Cellular Metabolic Landscape Identifies a Refractory Environment for Flavivirus Replication": Gullberg et al_Supplemental Figures

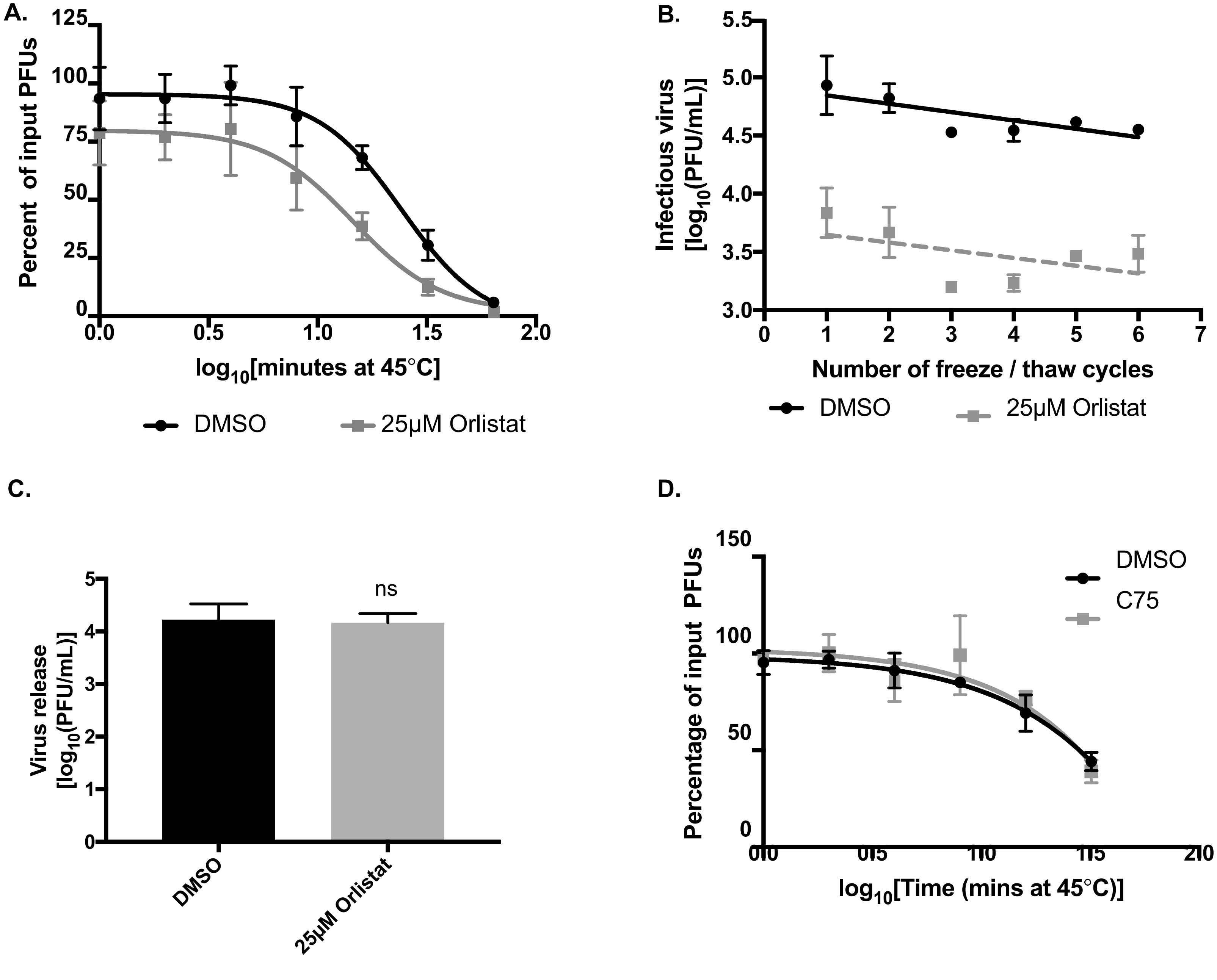


Gullberg et al, Supplemental Figure 1.

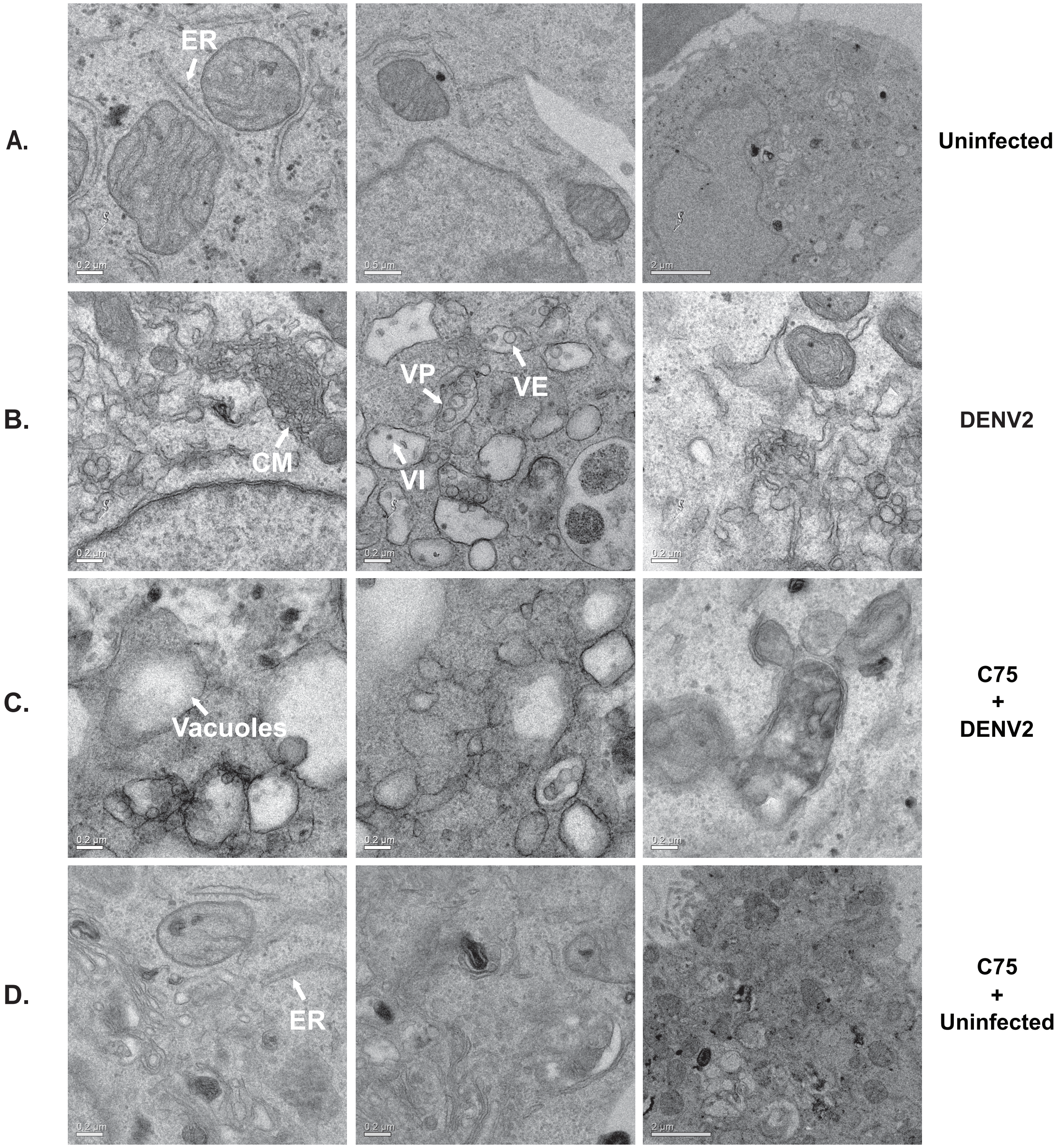


E.

| Treatment | Vesicles (per cell section) | CM (per cell section) |
| --- | --- | --- |
| **DENV2** | 37 | 0.62 |
| **C75 + DENV2** | 38 (No reduction) | 0.18 (70% reduction) |

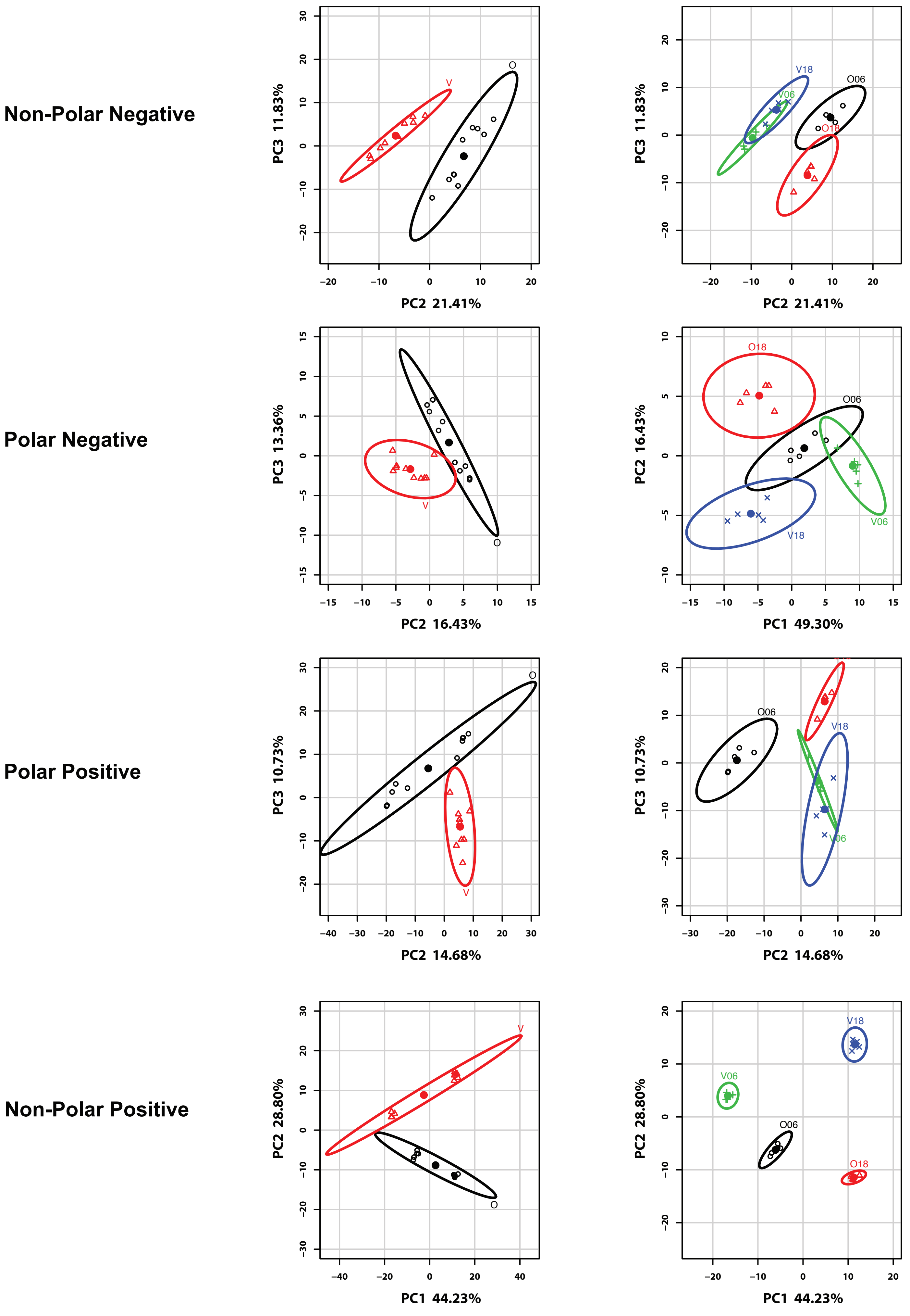


Gullberg et al, Supplemental Figure 3.

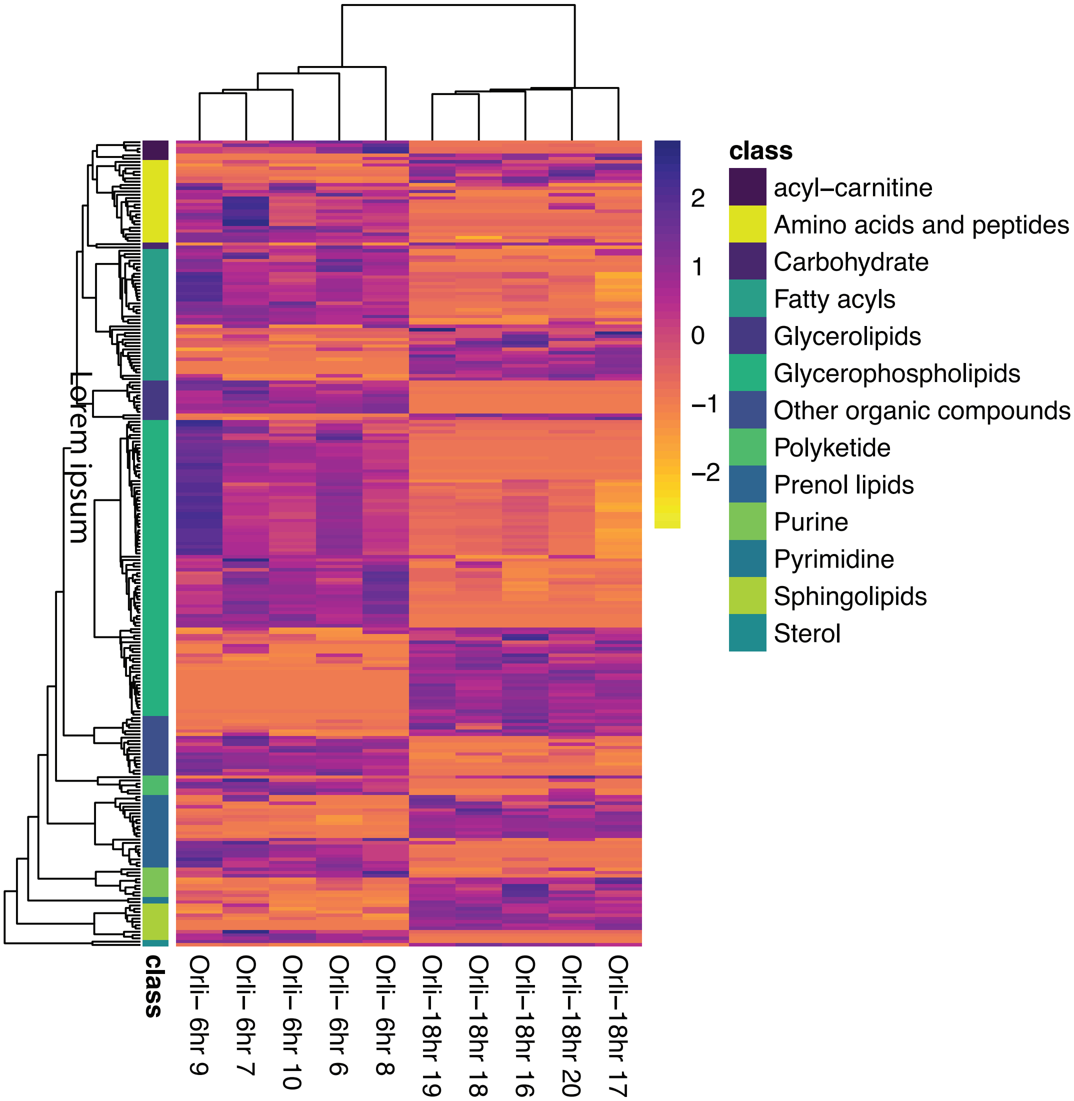


**Gullberg et al, Supplementary Figure 4**

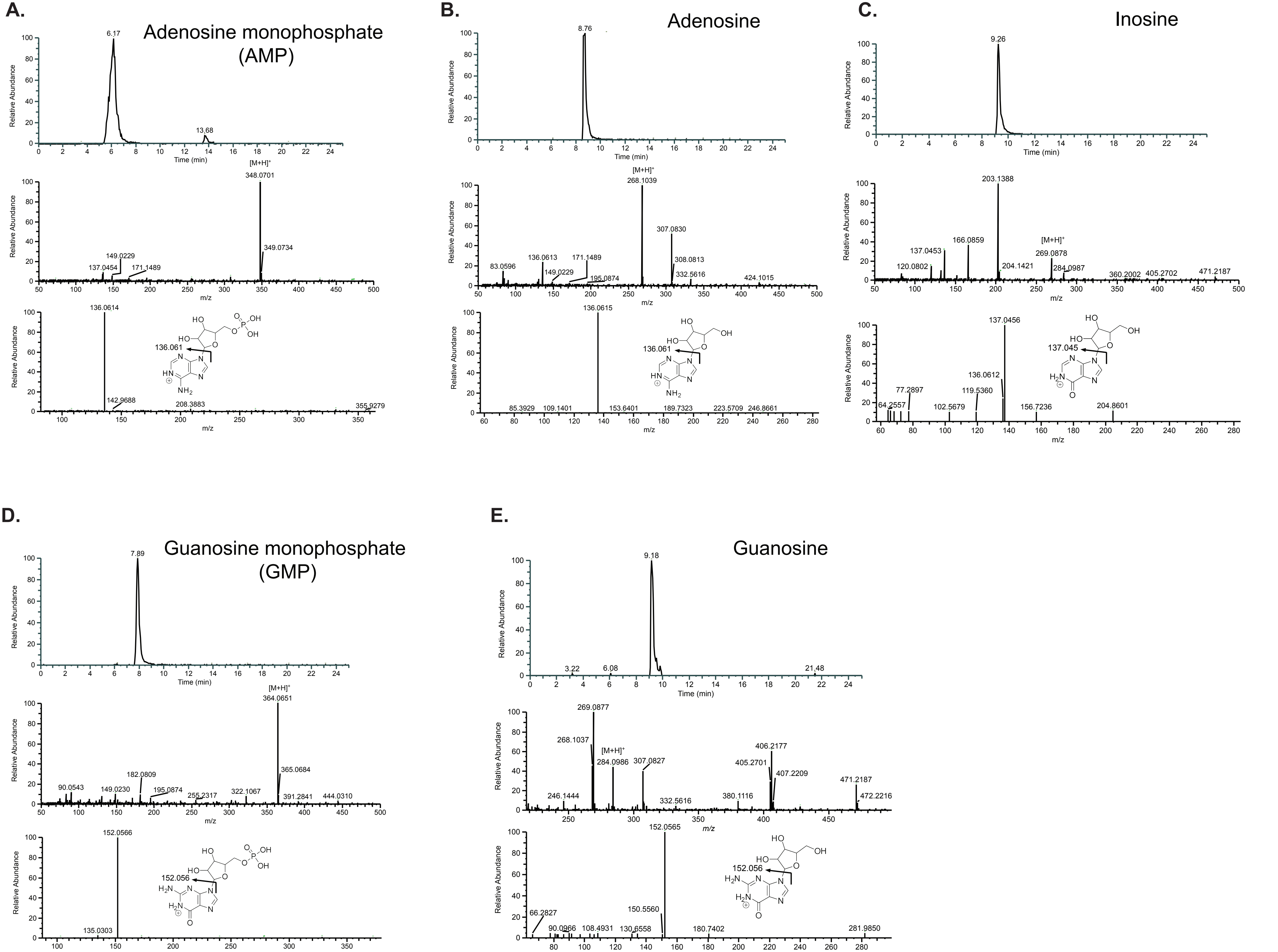


**Gullberg et al, Supplemental Figure 5.**
